# A unified model for microtubule rescue

**DOI:** 10.1101/401372

**Authors:** Colby P. Fees, Jeffrey K. Moore

## Abstract

How microtubules transition from depolymerization to polymerization, known as rescue, is poorly understood. Here we examine two models for rescue: 1) an ‘end-driven’ model in which the depolymerizing end stochastically switches to a stable state; and 2) a ‘lattice-driven’ model in which rescue-sites are integrated into the microtubule prior to depolymerization. We test these models using a combination of computational simulations and *in vitro* experiments with purified tubulin. Our findings support the ‘lattice-driven’ model by identifying repeated rescue sites in microtubules. In addition, we discover an important role for divalent cations in determining the frequency and location of rescue sites. We use ‘wash-in’ experiments to show that divalent cations inhibit rescue during depolymerization, but not during the polymerization. We propose a unified model in which rescues are driven by embedded rescue sites in microtubules, but the activity of these sites is influenced by changes in the depolymerizing ends.

## Introduction

Transitions between microtubule polymerization and depolymerization involve major structural rearrangements at the microtubule end. During polymerization, incoming tubulin subunits first form longitudinal interactions to generate curved protofilaments that extend from the microtubule end, and then form lateral interactions with neighboring protofilaments to straighten into the coherent microtubule lattice ^1^. During depolymerization, protofilaments undergo the opposite transition; from the straight conformation of the lattice, to losing lateral interactions and sharply curling away from each other at the disassembling end ^1–3^. Under some conditions, protofilaments at depolymerizing microtubule ends adopt a highly curled ‘ram’s horn’ morphology that is not observed for polymerizing microtubules ^2,3^. The outward curling of protofilaments during depolymerization is estimated to generate at least 19pN*nm of energy per tubulin subunit, and is thought to propagate strain along the protofilament that drives the straight-to-curled transition for subunits further down the lattice ^4^.

The transition from polymerization to depolymerization, known as catastrophe, must represent an intermediate state between the polymerizing and depolymerizing end structures. Current models posit that catastrophe may be caused by changes in the so-called ‘GTP cap’ at the growing plus end, which would weaken subunit affinity for the lattice ^5–7^. Alternatively, catastrophe may be caused by the formation of asymmetric protofilament extensions during polymerization that lack stabilizing lateral interactions and lead to a weakening of the lattice ^8,9^. Both of these models imply that a quorum of stable lateral interactions is necessary to maintain polymerization, and catastrophe may represent a loss of quorum.

In contrast to catastrophe, the transition from depolymerization to polymerization, known as rescue, is poorly understood. In principle, rescue must represent an intermediate state between the depolymerizing and polymerizing end structures. This intermediate would need to overcome both the fast rate of subunit loss and the energy driving the outward curling of protofilaments. Rescue is a property intrinsic to tubulin, as it can be observed in experiments using purified tubulin ^10–12^. Importantly, this work showed that rescue frequency is largely independent of tubulin concentration, indicating that the mechanism is not simply kinetic (i.e. the subunit on rate does not overwhelm the subunit off rate during depolymerization ^13,14^). Instead, rescue may disrupt the depolymerizing end structure. Several conditions have been shown to promote rescue, including mechanical stress on the microtubule ^15^, structural defects in the lattice ^12^, slowing GTP hydrolysis ^11^, and extrinsic regulation by microtubule-binding proteins ^9,16–21^. Nevertheless, we lack a basic understanding of the underlying mechanism.

A fundamental question is whether rescues are a pre-determined property of the microtubule lattice or a state that may be sampled by the depolymerizing end. In the first model, ‘rescue sites’ are formed in the lattice of a growing microtubule. After a subsequent catastrophe, when the depolymerizing end returns to a rescue site in the lattice, further depolymerization is inhibited and the microtubule end returns to a polymerizing state. This model is supported by several observations, including rescues near ‘GTP islands’ which are detected in microtubule lattices, far from polymerizing ends ^10^, and rescues near sites of mechanically induced damage, where subunits are thought to be lost from the lattice and new subunits may be incorporated ^12^. In the second model, changes in the structure and/or the activity of the depolymerizing microtubule end disrupt depolymerization and promote polymerization. This model is primarily supported by previous studies of proteins that selectively bind to microtubule ends and promote rescue, including CLASP, CLIP-170, and Kip3/kinesin 8 ^16,17,19–24^. It is unclear whether this is also a property of tubulin alone.

Here we tested these two models through a combined approach; first we developed Monte Carlo simulations to test simple predictions of each model. Then we validated the simulations experimentally using purified tubulin and found support for a combination of the two models. We find that regions of the microtubule lattice were capable of repeated rescues, which strongly supports the lattice-driven model. In addition, we observed that divalent cations are potent inhibitors of microtubule rescue, and used this to test whether conditions during lattice polymerization or depolymerization determine rescue activity. Our results support a combination of the two models wherein rescues are regulated through coordination between defined rescue sites in the lattice and changes at the depolymerizing end.

## Results

### Microtubule rescue frequency is independent of tubulin concentration

We sought to investigate the mechanism of microtubule rescue that is intrinsic to tubulin. Using purified tubulin to study microtubule rescue *in vitro* has been historically challenging, because rescues occur infrequently, compared to catastrophes. Therefore, we developed an experimental protocol to address three limiting factors of studying rescue *in vitro*; 1) temporal resolution, 2) acquisition duration and 3) microtubule number. Microtubules were assembled from purified porcine brain tubulin; determined to be >99% pure by Coomassie stain following SDS-PAGE separation (Figure 1A). This purified tubulin was mixed with 15-20% Hylite-488 labeled-tubulin to visualize microtubule polymerization from Rhodamine-labeled GMPCPP seeds using TIRF microscopy and standard polymerization conditions (5-10 µM tubulin in BRB80 with 5 mM MgCl_2_, 1 mM GTP and oxygen savaging buffer ^25,26^; Figure 1B). We imaged thousands of microtubules at a 1-second time resolution for 15 minutes using a cMOS camera and processed the images into kymographs to measure microtubule dynamics using a custom MATLAB program^25^. Measured rates of polymerization and depolymerization were used to determine the apparent tubulin association and dissociation constants, respectively (K_on_: 3.97 subunits*µM^−1^*s^−1^ K_off_: 567.30 subunits*s^−1^; Figure 1C & D). The catastrophe frequency (growing to shrinking) was calculated by dividing the number of transitions by the total polymerization time. We found no correlation between tubulin concentration and catastrophe frequency (Figure 1E). This experimental design allowed us to consistently detect rescues within the microtubule population and test models of the rescue mechanism.

**Figure 1.**
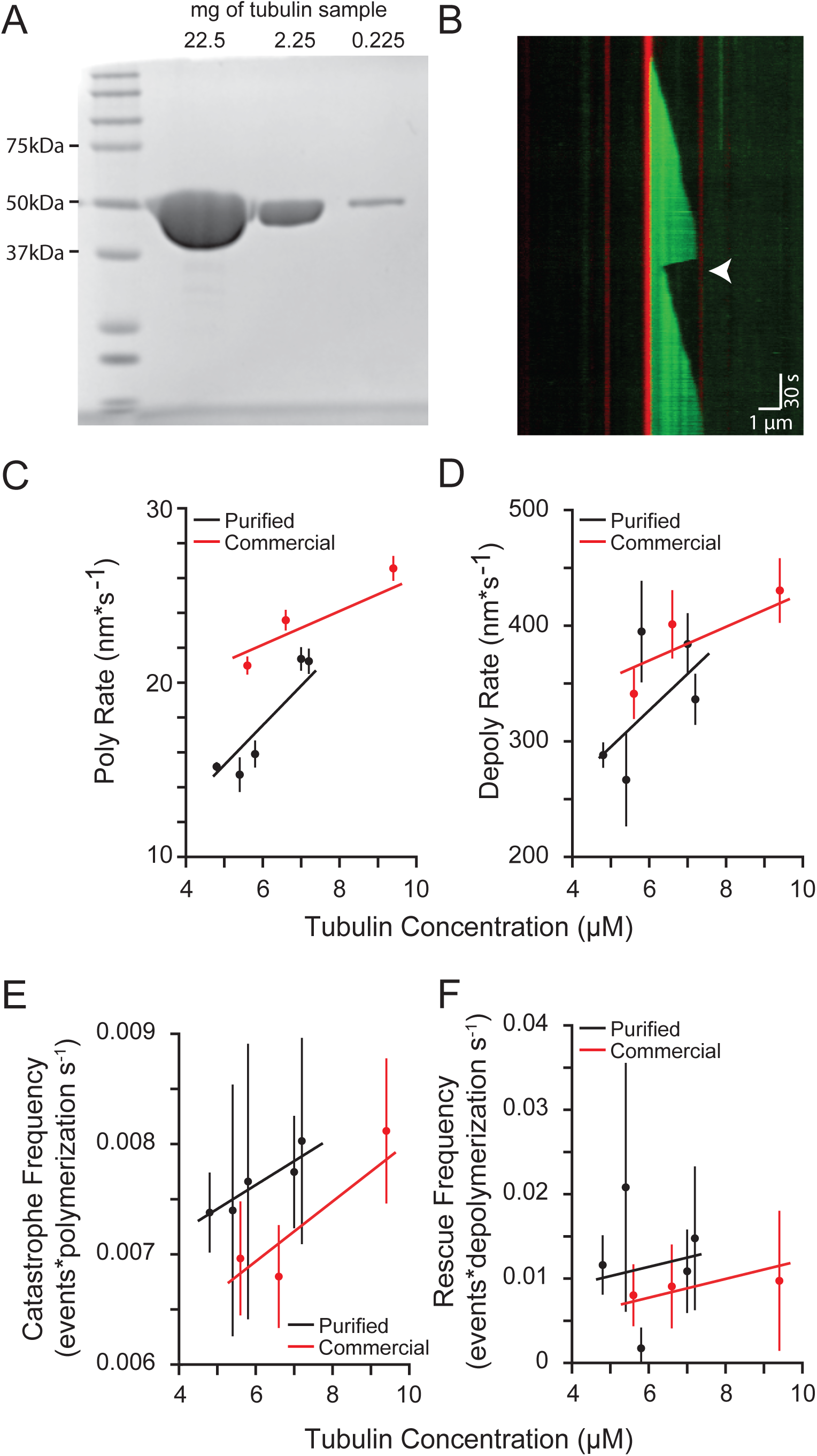
Microtubule rescue frequency is independent of tubulin concentration. **(A)** Purified tubulin sample separated by SDS-PAGE gel stained with Coomassie. Lane 1: protein ladder. Lanes 2-4: 10-fold serial dilutions of tubulin purified from Porcine brain. Protein loads are approximately 22.5 – 0.225 mg, respectively. **(B)** Representative kymograph of purified tubulin (green) polymerized from GMPCPP stabilized microtubule seeds (red), rescue event denoted (arrowhead). Images were collected at 1-second intervals. Tubulin concentration: 7.2 µM, vertical scale bar = 0.5 minute and horizontal scale bar = 1 µm. **(C)** Polymerization rate plotted as a function of tubulin concentration. Protein purified from pig brain (black) compared to purchased protein purified commercially (red). Data points are mean ± 95% CI. **(D)** Depolymerization rate plotted as a function of tubulin concentration. **(E)** Catastrophe frequency plotted as a function of tubulin concentration. Catastrophe frequency defined as the number of events per sum of polymerization time for each concentration. Data points represent the mean of catastrophe frequencies measured per microtubule, from all microtubules at a given tubulin concentration. Error bars represent 95% CI. **(F)** Rescue frequency plotted as a function of tubulin concentration. Rescue frequency calculated as the number of rescues divided by total depolymerization time for each concentration. Data points represent the mean value of rescue frequencies measured per microtubule, from all microtubules at a given tubulin concentration. Error bars represent 95% CI. For figures 1C-F, at least 73 microtubules were analyzed for each tubulin sample, at each concentration.

We defined rescue as a transition from depolymerization to polymerization that occurred at least 150 nm from the GMPCPP-stabilized seed (Figure 1B, arrowhead). This definition ensures the distinction of rescues that occur in the dynamic microtubule lattice from those that occur at the stabilized seed, within the temporal and spatial resolution of our system. We measured rescues across a range of tubulin concentrations and found an average rescue frequency of 0.01 events per second of depolymerization time (CI: 0.009 – 0.014 events*s^−1^), with no clear dependence on tubulin concentration (Figure 1F). Overall, 18.7% of the 525 total seeds in our experiments nucleated microtubules that exhibited at least one rescue, and these rescued an average of 1.6 times per 15-minute acquisition (CI: 1.4-1.7; Supplemental Figures 1A & 1B).

We then validated the results from our purified porcine brain tubulin by conducting parallel experiments with commercially available porcine brain tubulin under the same reaction conditions. The commercially available tubulin exhibits similar polymerization and depolymerization rates as our purified tubulin (Figures 1C & D), and similar frequencies of catastrophe and rescue (Figures 1E & 1F). Importantly, neither set of experiments revealed significant correlation between tubulin concentration, and therefore polymerization rate, and rescue frequency, consistent with previous findings ^13,27^.

### Longer microtubules are more likely to rescue

We considered two general models for how rescues might be triggered. First, an ‘end-driven’ model in which the depolymerizing end of the microtubule stochastically switches to a stable state, resulting in rescue. Second, we considered a ‘lattice-driven’ model in which rescuesites were integrated into the lattice prior to depolymerization and produced a rescue when the depolymerizing end reached that site in the lattice. Both models predict that longer microtubules would have a greater likelihood of rescue. Consistent with our prediction, we found that microtubules that rescue in our experiments were 51% longer at catastrophe than the population on average (mean = 3.68 µm, CI: 3.25 – 4.11 µm, compared to 2.44 µm, CI: 2.32 – 2.55 µm; Figure 2A).

**Figure 2.**
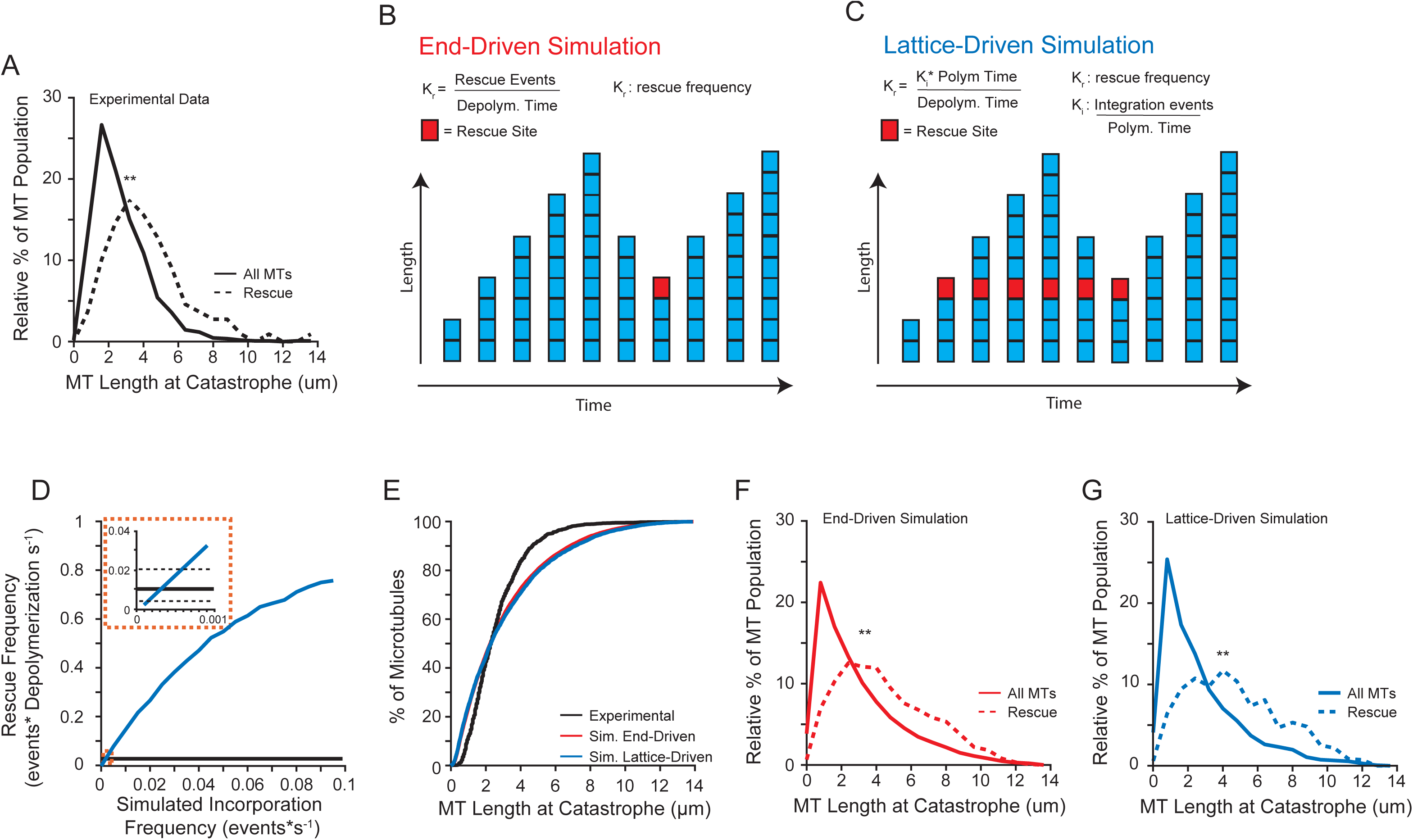
Longer microtubules are more likely to rescue. **(A)** Relative distribution of microtubule lengths at catastrophe leading to rescue (dashed line) compared to the entire population (solid line) of the experimental data under standard conditions (5mM MgCl_2_). Data represents 1127 microtubule lengths from 525 stabilized seeds, pooled from 5 separate experiments. **(B)** Cartoon schematic of our end-driven simulation. Rescue site is denoted by a red square. Rescue frequency is defined as the number rescue events per depolymerization time. Rescues were triggered stochastically during depolymerization at the experimentally derived rescue frequency (0.01 events*depolymerization s^−1^). **(C)** Cartoon schematic of our lattice-driven simulation. Rescue sites (denoted by a red square) are incorporated during polymerization, and always trigger rescue when the depolymerizing end returns to that site. **(D)** Simulated rescue frequency as a function of incorporation frequency. The simulated rescue frequency for each incorporation rate calculated from 1000 simulated microtubules. **(E)** Cumulative distribution of microtubule lengths at catastrophe under standard conditions (black) compared to end-driven simulation (red) and the lattice-driven simulation (blue). **(F)** Similar to figure 2A, using data from the end-driven simulation. Data represents 60,000 simulated microtubules. **(G)** Similar to figure 2A, using data from the lattice-driven simulation, with the incorporation frequency set to 0.0001 events*s^−1^. Plot represents 60,000 simulated microtubules.

We then developed computational models of microtubule rescue using Monte Carlo simulations of experimentally derived parameters (polymerization and depolymerization rates as well as transition frequencies; Table 1) to simulate a single filament with changes in length over time as the output. For our ‘end-driven’ simulation, rescue events were stochastically triggered during depolymerization, at a rescue frequency (K_r_) that matched our experimental measurements (Figure 2B). For our ‘lattice-driven’ simulation, rescue sites were incorporated during filament polymerization at a defined frequency (K_i_). These sites would remain embedded in the lattice until, after catastrophe, the depolymerizing end reached the site and triggered a rescue event. Initially we set these sites as fully active – if they were incorporated into a filament, then that site would always trigger a rescue, however this was modified later (see below; Figure 2C). The K_i_ was computationally determined using our simulation to calculate the rescue frequency as a function of incorporation frequency (Figure 2D). The simulated rescue frequency intercepted the experimental values at an incorporation frequency between 0.0001 – 0.0006 events*s^−1^ (Figure 2D inset).

**Table 1:** Common Simulation Parameters

| Parameter | Value |
| --- | --- |
| Nucleation sites | 1 |
| Nucleation Rate | $0.02 \text{ s}^{-1}$ |
| Time Resolution | 1 s |
| Duration | 15 min |
| Microtubules per Simulation | 20,000 |
| Total Tubulin | 5-8 $\mu\text{M}$ |
| $K_{\text{on}}$ (subunits*$\mu\text{M}^{-1}\text{s}^{-1}$) | 3.97 |
| $K_{\text{off}}$ (subunits*$\text{s}^{-1}$) | 567.30 |
| $V_g$ (nm*$\text{s}^{-1}$) | 17.77 (17.28 – 18.27) |
| $V_s$ (nm*$\text{s}^{-1}$) | 324.16 (313.65 – 334.69) |
| $K_c$ ($\text{s}^{-1}$) | 0.0029 (0.0026 – 0.0032) |
| $K_R$ ($\text{s}^{-1}$) | 0.012 (0.009 – 0.014) |
| $K_I$ ($\text{s}^{-1}$)<sup>§</sup> | 0.001 |

§: Applies to Lattice-driven Model.

**Table 2:** Model Simulation Results

| Parameter | Lattice-Driven Model | End-Driven Model | Experimental |
| --- | --- | --- | --- |
| $K_R$ ($\text{s}^{-1}$) | 0.007 (0.006 – 0.007) | 0.010 (0.001 – 0.011) | 0.012 (0.009 – 0.014) |
| $V_s$ (nm*$\text{s}^{-1}$) | 301.23 (300.15 – 302.30) | 303.42 (303.87 – 306.96) | 324.16 (313.65 – 334.69) |
| MT<sub>Length</sub> ($\mu\text{m}$) | 3.0 (2.9 – 3.0) | 3.1 (3.0 – 3.1) | 2.8 (2.7 – 2.9) |
| % Repeated Rescues | 0.18 | 0.49 | 35 |

We validated our end-driven and lattice-driven simulations first by comparing the lengths of microtubules at catastrophe to our experimental data. We found that both simulations accurately sample a similar range of microtubule lengths at catastrophe (Figure 2E). Of note, the simulated data was more normally distributed for microtubules that catastrophe above 6 µm in length. We attributed this to a limitation of our experimental setup, specifically the inability to monitor longer microtubules that extend beyond the field of view. Both simulations were consistent with our experimental results showing that rescuing microtubules were significantly longer at catastrophe than the average distribution of lengths (Figures 2F & 2G). We then sought to use these simulations to determine the mechanism of microtubule rescue by testing predictions from each model.

### Effect of depolymerization rate on rescue

We began by testing whether depolymerization rates influenced rescue frequency. If rescue occurs through stochastic transitions at the depolymerizing end, then increasing the time spent in the depolymerizing state might promote rescue. We used our simulations to test this prediction across a range of experimentally derived depolymerization rates (Figure 3A). The end-driven simulation showed that rescue events were preceded by significantly slower depolymerization rates (mean = 250.7, CI: 240.3-261.1 nm*s^−1^) than the rates exhibited by the total population (288.1, CI: 283.8-292.3 nm*s^−1^; Figure 3B). In contrast, the lattice-driven simulation showed no difference in depolymerization rates preceding a rescue event compared to the total population (Figure 3C). Our experimental data showed no difference in the depolymerization rates preceding a rescue event compared to the total population (mean = 274.5, CI: 243.73 – 305.3 nm*s^−1^, compared to 289.8, CI: 276.7-302.8 nm*s^−1^; Figure 3D). In this case, our experimental data is more consistent with the lattice-driven model.

**Figure 3.**
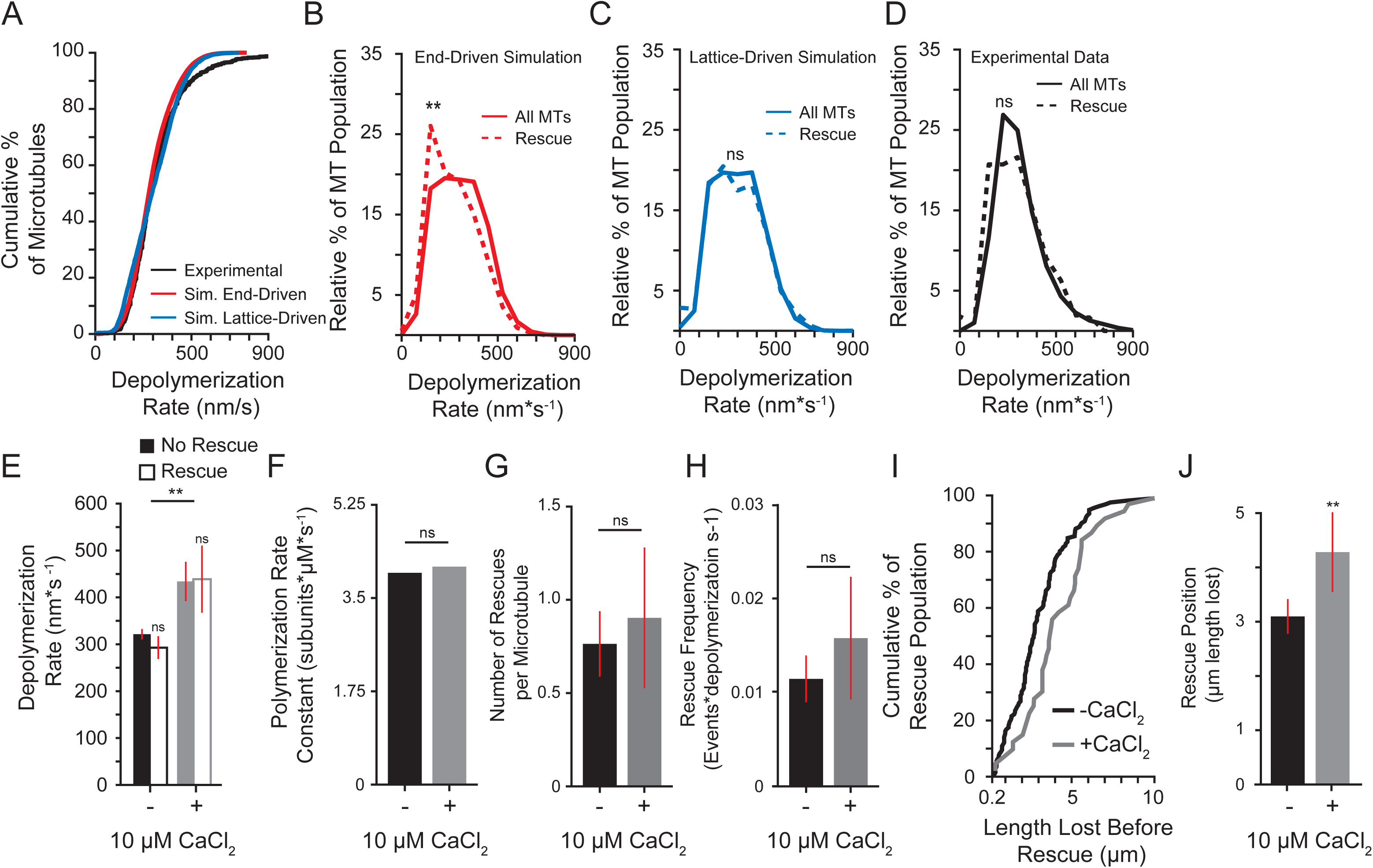
Effect of depolymerization rate on rescue. **(A)** Cumulative distribution of depolymerization rates of experimental data under standard conditions (5 mM MgCl_2_; black) compared to the end-driven simulation (red) and the lattice-driven simulation (blue). **(B)** Relative distribution of depolymerization rates leading to rescue (dashed line) compared to entire population (solid line) of the end-driven simulation. Plot represents 60,000 simulated microtubules. **(C)** Relative distribution of depolymerization rates leading to rescue (dashed line) compared to entire population (solid line) of the lattice-driven model. **(D)** Relative distribution of depolymerization rates leading to rescue (dashed line) compared to entire population (solid line) from experimental data under standard conditions (5 mM MgCl_2_). Rescue data represents 165 depolymerization events from 112 microtubules; solid line represents entire population data consisting of 1283 depolymerization events from 1164 microtubules from 5 separate experiments. **(E)** Average depolymerization rates from experimental data of catastrophes terminating at the seed (No Rescue) compared to those terminating at a rescue site within the lattice (Rescue) with and without 10 µM CaCl_2_ (Black and grey respectively). Significance determined by comparing rescue to no rescue for each condition as well as rescue across conditions. Bars represent mean ± 95% CI, at least 230 depolymerization events from at least 3 experiments were examined for each condition. **(F)** Average polymerization rate constants from experimental observations of at least 275 microtubules for each condition. **(G)** Average number of rescues per microtubule with and without 10 µM CaCl_2_. Bars represent mean ± 95% CI from at least 275 microtubules over at least 3 separate experiments for each condition. **(H)** Average rescue frequency calculated as events per depolymerization time. Bars represent mean ± 95% CI from at least 275 microtubules over at least 3 experiments per condition. **(I)** Cumulative distribution of the length lost before rescue with and without CaCl_2_. **(J)** Mean ± 95% CI of the rescue position relative to the catastrophe site. Position defined as the difference of microtubule length at catastrophe and length at rescue. Bars represent at least 39 length measurements from 25 microtubules from at least 3 separate experiments. ** p << 0.001, ns: not significant. Significance determined by Mann-Whitney U test.

The results from our end-driven simulation showed that the average difference in depolymerization rates preceding rescue was approximately 30 nm*s^−1^, which is at the edge of the detection threshold of our experiment. As an alternative approach, we tested the prediction that decreasing the time spent in depolymerization, by increasing the rate of depolymerization, would inhibit rescue. The divalent cation calcium (Ca^2+^) increases microtubule depolymerization rates^28–31^ without significantly altering polymerization rates^29^. Adding 10 µM CaCl_2_ to our experimental conditions significantly increased the depolymerization rate compared to control reactions (+CaCl_2_: mean = 442.0, CI: 403.3 – 480.8 nm*s^−1^ compared to no CaCl_2_: 324.4, CI: 314.2 – 334.9 nm*s^−1^; Figure 3E), without altering the apparent polymerization rate constant (Figure 3F and Supplemental Figure 2). Under these conditions, we did not observe a significant change in the number of rescues per microtubule or the rescue frequency as a function of depolymerization time (Figure 3G & H). However, we observed that more lattice was lost during depolymerization in the presence of 10µM CaCl_2_ before rescue occurred (Figure 3I). We calculated rescue position relative to catastrophe position as the difference of microtubule length at catastrophe and length at rescue, which we will refer to as rescue position. We found that adding CaCl_2_ significantly shifted the position of rescue sites, causing more lattice loss before rescue (Figure 3J). These results suggest that depolymerization rate influences the position, but not the timing of rescue.

### Rescues occur repeatedly at similar sites along the microtubule

How might the position of rescue sites be determined? We predicted that if rescues were triggered by pre-defined sites integrated into the lattice, then the same sites might exhibit multiple rescues in our experiments. Consistent with this prediction, we observed 35% of rescues occur within the same region (± 200 nm) as a prior rescue (Figure 4A arrowheads). We termed these events ‘repeated rescues’.

**Figure 4.**
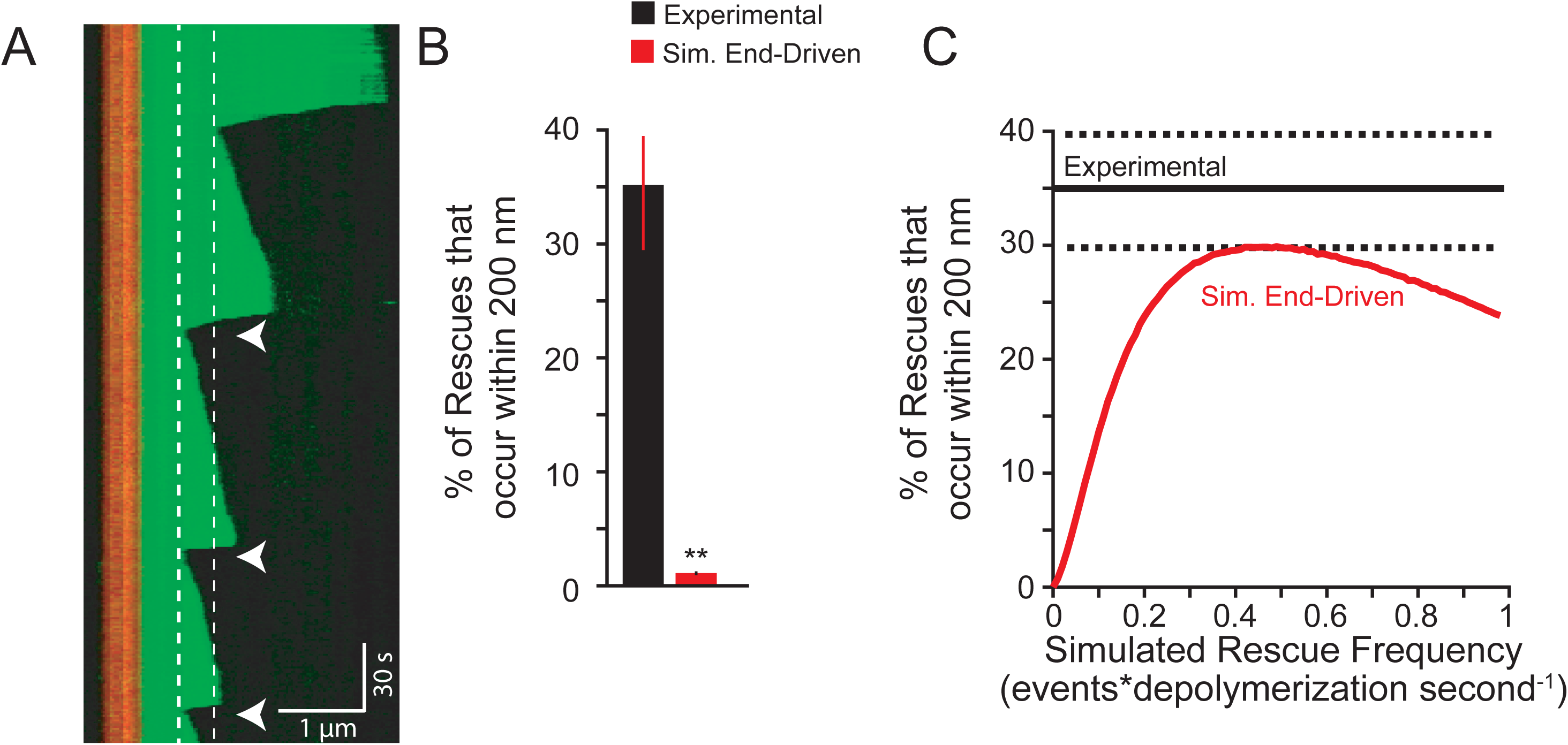
Rescues occur repeatedly at similar sites along the microtubule. **(A)** Representative kymograph showing repeated rescue events (within 200 nm of previous rescue denoted with arrowheads. Dashed lines highlight the location of two separate rescue sites through time. Images were collected at 1-second intervals. Tubulin concentration: 5.6 µM, vertical scale bar: 30 s, horizontal scale bar: 1 µm. **(B)** Percentage of the rescue population that rescues repeatedly (within 200nm) under standard conditions (black) compared to simulated results of the end-driven simulation (red). Experimental data represented as mean ± SEP, pooled from 105 total rescues events from 5 separate experiments. Simulated data represents 60,000 microtubules from 10 separate simulations. **(C)** Percent of repeated rescues calculated with increasing rescue frequencies using the end-driven simulation (solid red line). Experimental data represented as mean (solid black line) ± SEP (dashed black lines). Sets of simulations were preformed using 1,000 microtubules per rescue frequency tested. Rescue frequencies were increased by 0.005 intervals between 0.01-0.99 events*s^−1^. Each simulation set was then repeated 5 separate times. Simulated data was pooled and plotted as mean ± SEP for each rescue frequency. ** p << 0.001. Significance determined by Fishers exact test.

We used our simulation to test whether repeated rescues could be generated by an end-driven mechanism alone. Starting with our experimentally derived rescue frequency (0.01 events per second of depolymerization time), our end-driven simulation showed that <0.5% of rescues occurred within 200nm of a previous rescue site (Figure 4B red bar). We then asked what rescue frequency would be required to generate the repeated rescue behavior observed in our experiments? We ran multiple end-driven simulations and found that the overall rescue frequency must be ∼40x greater (i.e. 0.4 events per second of depolymerization time) to generate repeated rescue behavior that matches our experimental results (Figure 4C). This suggests that an end-driven model alone is highly unlikely to generate repeated rescues. The lattice is likely to play a determining role.

### Divalent cations suppress rescues

We next sought to identify factors that influence rescue sites. Recent work by Aumeier et al. reported microtubule rescues occurring closer to the site of catastrophe than we observed in our standard experimental conditions ^12^. An important difference between their experiments and our own is the concentration of magnesium – whereas Aumeier et al. used polymerization buffer containing 1mM MgCl_2_, our polymerization buffer contains 5mM MgCl_2_ ^12^. We confirmed the impact of magnesium on rescue by altering MgCl_2_ concentration in our experiments. Rescues occur closer to the site of catastrophe in 1mM MgCl_2_ compared to 5mM MgCl_2_ (1mM Mg: mean = 1.4, CI: 1.4 – 1.7 µm; 5mM Mg: 3.0, CI: 2.7 – 3.3 µm; Figure 5A and B). In other words, lower magnesium decreases the amount of the microtubule lattice that depolymerizes before rescue occurs. We further tested the impact of magnesium by titrating a range of MgCl_2_ concentrations in the reaction buffer. Consistent with previous studies, we found that rates of polymerization and depolymerization increase in proportion with the concentration of MgCl_2_ in the reaction (Supplemental Figures 3A & 3B; ^25,32^). Increasing MgCl_2_ concentration inhibits rescue, as a function of both rescue position relative to the site of catastrophe and time in depolymerization (Figures 5B & C). Adding 10µM CaCl_2_ also causes rescues to occur further from the site of catastrophe, even under low magnesium conditions (1mM MgCl_2_ + 10µM CaCl_2_; Figure 5A and 5B); however, adding 10µM CaCl_2_ did not increase the rescue frequency as a function of depolymerization time (Figure 5C). This is likely attributable to differences in depolymerization rate, as 10µM CaCl_2_ increases depolymerization rate by > 150% (Figure 3F). These results suggest that divalent cations strongly influence the position of rescue sites.

**Figure 5.**
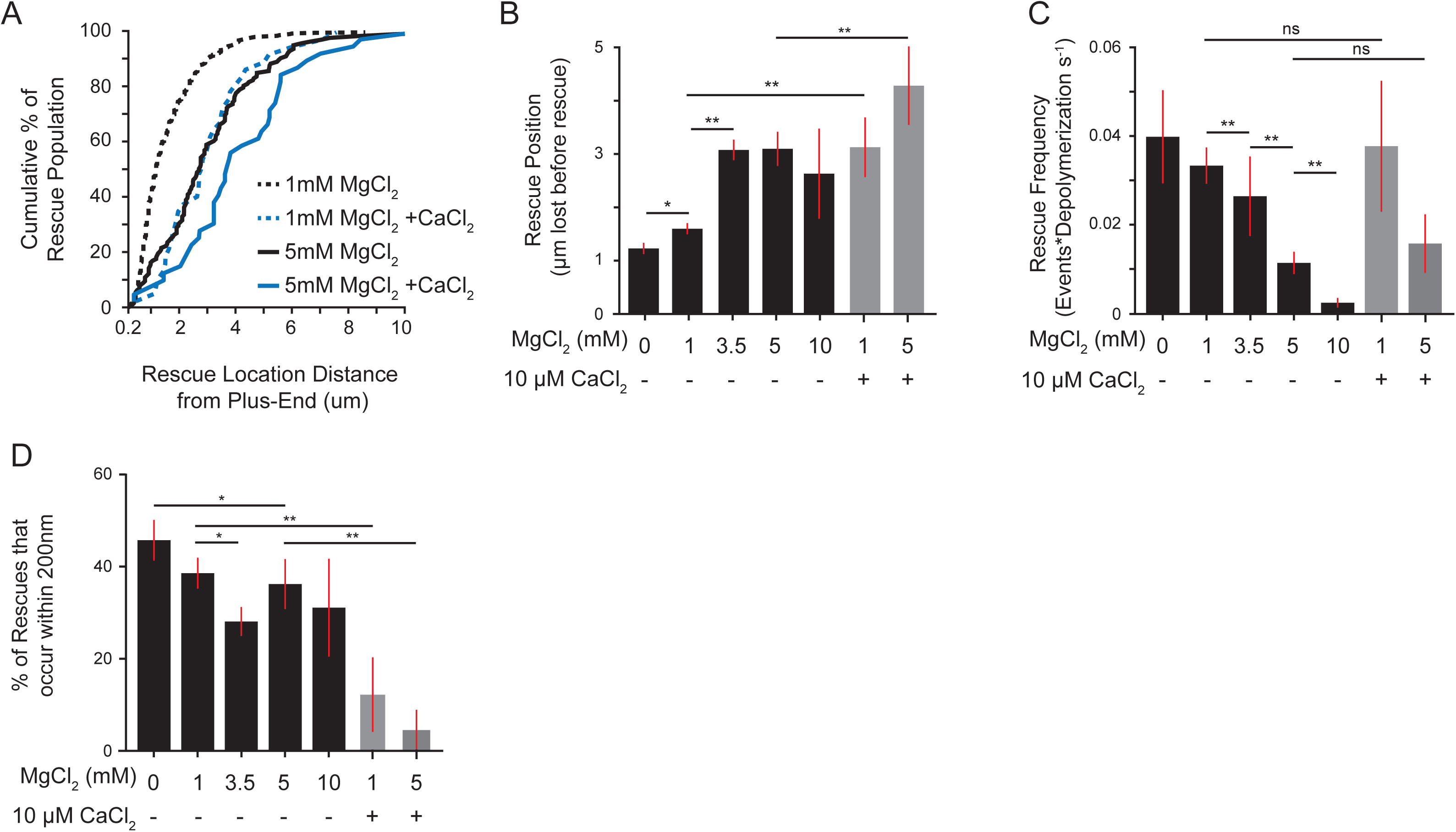
Divalent cations suppress rescues. **(A)** Cumulative distribution of microtubule length lost before rescue. Solid lines are 5mM MgCl_2_ without (black) or with 10µM CaCl_2_ (blue). Dashed lines are 1mM MgCl_2_ without (black) or with 10µM CaCl_2_ (blue). Each line represents at least 39 lengths from at least 25 microtubules pooled from at least 3 separate experiments. **(B)** Rescue position relative to catastrophe site. Bars represent mean ± 95% CI from at least 39 length measurements from 25 microtubules, pooled from at least 3 separate experiments. **(C)** Rescue frequency as a function of depolymerization time. Bars represent mean ± SEP for each condition. Data pooled from at least 87 microtubules from at least 3 separate experiments for each condition. **(D)** Percent of rescues that occur repeatedly, as in figure 4B. Bars represent mean ± SEP. Figure 5E: significance determined by Fishers exact test. All other panels: significance determined by Mann-Whitney U test. *: p < 0.01, **: p << 0.001, ns: not significant.

We then asked whether divalent cations alter repeated rescue activity in the lattice. We observed that increasing concentrations of MgCl_2_ decreases the frequency of repeated rescues (45% of rescues at 0mM MgCl_2,_ compared to 25% of rescues at 3.5mM MgCl_2_ Figure 5E). Furthermore, adding 10 µM CaCl_2_ strongly decreases the frequency of repeated rescues (Figure 5E). Together these results indicate that divalent cations suppress rescue.

### Rescue activity is determined during depolymerization

We reasoned that divalent cations could suppress rescue either by inhibiting the integration of rescue sites into the lattice or inhibiting their activity. To distinguish between these possibilities, we asked whether divalent cations influence rescues during the polymerization of the lattice or during depolymerization. We designed an experiment to isolate lattice polymerization from depolymerization by assembling microtubules under one set of reaction conditions, then rapidly exchanging the reaction buffer and testing how the pre-formed lattice behaves under different conditions (Figure 6A). These experiments differ from previously described ‘wash-out’ experiments, in which the free tubulin was depleted from the reaction to halt further polymerization ^14,25,32^. In contrast, our ‘wash-in’ experiments maintain the same concentrations of tubulin, GTP, and oxygen scavenging buffer throughout the experiment; only the concentration of MgCl_2_ is changed. Accordingly, we observed that after buffer exchange, microtubules continued to polymerize from the existing lattice at rates consistent with the present MgCl_2_ concentration. Increasing or decreasing concentrations of MgCl_2_ was sufficient to increase or decrease the polymerization and depolymerization rates, respectively (Figures 6B and Supplemental Figures 3A & 3B). Control experiments where reaction buffers with the same concentration of MgCl_2_ were washed in exhibited no difference in either polymerization or depolymerization rates (Figures 6B & 6C).

**Figure 6.**
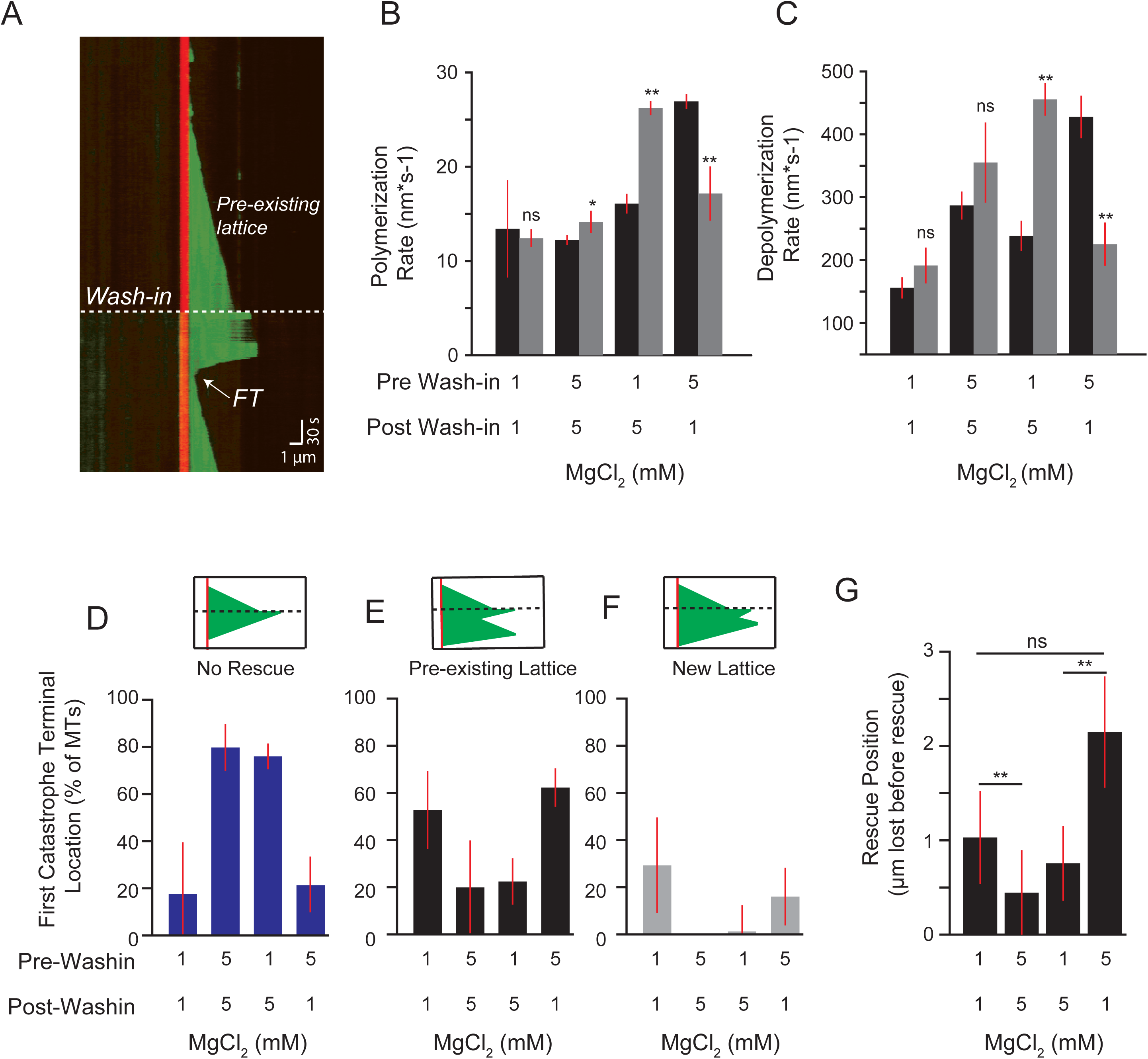
Rescue activity is determined during depolymerization. **(A)** Representative kymograph of wash-in experiment. Dashed line denotes the wash-in timepoint. Dashed line denotes the pre-existing lattice prior to wash-in. The first termination site after wash-in (FT) denoted with arrow (see Figures 6D-F). Image acquisition was paused during wash-in and resumed immediately afterwards (see Materials and Methods). Images were collected at 1-second intervals. Tubulin concentration: 3.2 µM, Prewash-in = 5 mM MgCl_2_ & post wash-in = 5 mM MgCl_2_, vertical scale bar = 30s & horizontal scale bar = 1 µm. **(B)** Polymerization rates of microtubules before (black bars) and after (gray bars) wash-in for each condition. Bars represent mean ± 95% CI of pooled data from at least 43 polymerization rates from 30 microtubules and a minimum of 3 separate experiments. Significance denotes differences between rates before and after wash-in. **(C)** Depolymerization rates of microtubules before (black bars) and after (gray bars) wash-in for each condition. **(D)** Percent of first catastrophes after wash-in that depolymerize to the stabilized seed and did not rescue. Bars in 6D – 6F represent mean ± SEP**. (E)** Percent of first catastrophes after wash-in that rescue within the pre-existing lattice. **(F)** Percent of first catastrophes after wash-in that rescue within lattice added after wash-in. **(G)** Mean position of the first rescue after wash-in. Bars represent mean ± 95% CI of data from at least 9 rescues events immediately after wash-in pooled from 3 separate experiments per condition. **: p << 0.001, ns: not significant. Significance determined by Mann-Whitney U test.

Using this experiment, we tested the prediction that if rescue is determined during the polymerization of the lattice, then microtubules grown under rescue-promoting conditions (1mM MgCl_2_) would be insensitive to the introduction of rescue-suppressing conditions (5mM MgCl_2_), and vice-versa. We tested this prediction by examining the first catastrophe event after wash-in and asked whether and where along the lattice the microtubules rescue. Outcomes were classified into one of three groups; 1) No rescue, in which depolymerization reached the stabilized seed (Figure 6D); 2) Rescue in pre-existing lattice, in which the microtubule rescued at a region of the lattice that was assembled prior to wash-in (Figure 6E); and 3) Rescue in new lattice, in which the microtubule rescued at a region of the lattice that was assembled after the wash-in (Figure 6F). Microtubules assembled in 1mM MgCl_2_ conditions (rescue-promoting) and then introduced to 5mM MgCl_2_ conditions (rescue-inhibiting) exhibited fewer rescues than controls that maintained 1mM MgCl_2_ (Figure 6D). In contrast, microtubules assembled in 5mM MgCl_2_ and then introduced into 1mM MgCl_2_ exhibited more rescues than controls that maintained 5mM MgCl_2_ (Figure 6D). Importantly, the majority of rescues after the introduction of 1mM MgCl_2_ occurred in regions of the lattice that were assembled prior to wash-in, in the presence of 5mM MgCl_2_ (Figure 6E and F). We also measured rescue position and found that introducing 1mM MgCl_2_ resulted in an average length of lattice lost before rescue that was similar to what we observed for microtubules that maintained 1mM MgCl_2_ throughout the experiment (Figure 6G). Together these results suggest that rescue activity is not exclusively determined during the polymerization of the lattice.

## Discussion

In this study, we sought to elucidate the mechanisms of microtubule rescue by testing two models: 1) a lattice-driven model, in which rescue-sites were integrated into the lattice, prior to depolymerization, and produced a rescue when the depolymerizing end returned to that site in the lattice; and 2) an ‘end-driven’ model, in which the depolymerizing end of the microtubule stochastically switches to a stable state, resulting in rescue. Our data are consistent with rescue sites integrating into the lattice, supporting the lattice-driven model; however, we also found that changing the depolymerization conditions after lattice polymerization, by altering the concentration of divalent cations in the reaction buffer, was sufficient to alter rescue behavior. We therefore propose a unified model, in which the lattice contains defined sites that can promote rescue, but the activity of these sites may be regulated by conditions during depolymerization.

Figure 7 depicts models for how features of the lattice and depolymerizing end could stabilize an intermediate state of the microtubule end, between depolymerization and polymerization. We speculate that the depolymerizing microtubule end is a heterogenous structure where individual protofilaments can visit different states of interactions with neighbors, curvature, and strain. In this model, states that enhance lateral interactions, reduce curvature, and/or reduce strain could lead to rescue. In addition, sites in the lattice could promote rescue by blocking the propagation of strain from the outwardly curled protofilament to the lattice. In this way, heterogenous regions of existing microtubule lattice disrupt depolymerization and promote the return to a polymerizing state.

**Figure 7.**
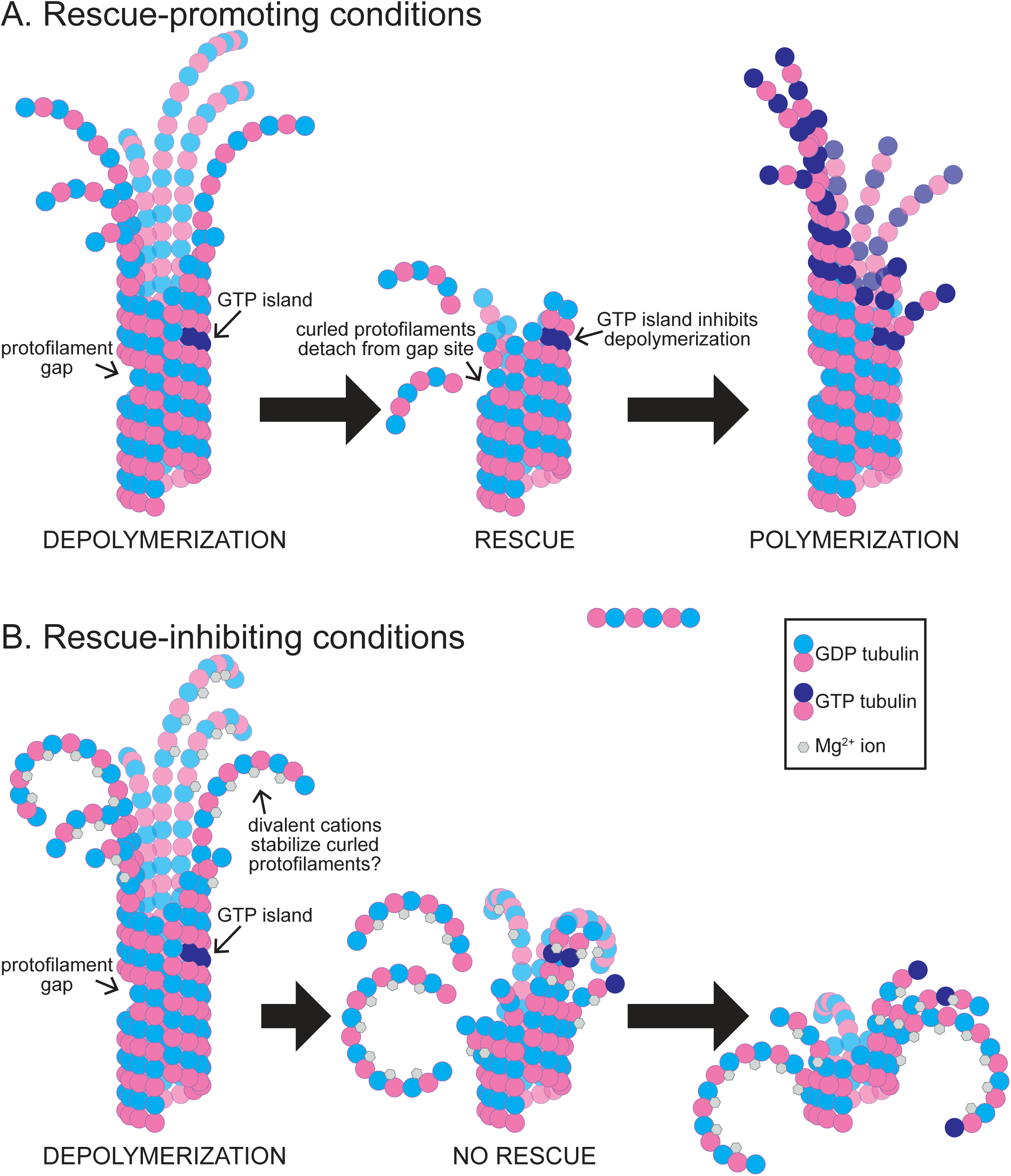
Rescue activity is determined during depolymerization. **(A)** Under rescue-promoting conditions, depolymerization is inhibited by rescue sites in the microtubule lattice, and the microtubule end returns to a polymerizing state. These sites may include, but are not limited to, regions of missing subunits (i.e. ‘protofilament gaps’) and subunits that maintain GTP at the E-site (i.e. ‘GTP islands’). **(B)** Under rescue-inhibiting conditions, divalent cations promote depolymerization and inhibit the activity of rescue sites in the lattice. Here we depict a proposed role for magnesium ions in promoting the outward curling of protofilaments.

The lattice-driven model is supported by several lines of evidence from previous studies, including the observation of apparent ‘GTP-islands’ -- regions of the lattice distal from the plus end that bind a GTP-tubulin specific antibody ^10^. These islands were shown to correlate with sites of rescue, suggesting that regions of incomplete GTP hydrolysis may stabilize the microtubule lattice to promote rescue ^10^. Along these lines, Tropini et al. demonstrated that adding small amounts of a non-hydrolysable GTP analog (GMPCPP) to microtubules assembling in GTP could increase the frequency of rescues^11^. Finally, recent work has shown that rescue events correlate with regions of microtubule repair, where new GTP-tubulin incorporates into sites along the lattice following mechanical damage ^12^. Our strongest evidence in support of the lattice-driven model comes from our observation of repeated rescue sites, which show successive rescues within the same region of lattice (Figure 4). Our simulations indicate that such behavior is not likely to arise from a stochastic rescue mechanism; therefore, the lattice at the repeated rescue site possesses a persistent, rescue-promoting activity. The exact nature of these repeated rescue sites in our experiments is uncertain, but clearly does not require the addition of non-hydrolysable nucleotide analogues or the induction of mechanical damage to microtubules. Those conditions may increase rescues, but they are not necessary for rescues to occur.

An important question that has not been addressed by previous studies is how long rescue sites remain active and what are the requirements for activity. In an effort to measure the lifetime of repeated rescue sites, we attempted to quantify the period of time over which repeated rescues can occur; however, we were only able to capture the complete lifetime of a few microtubules with repeated rescue sites. While sampling issues limit our ability to make general conclusions about the lifetime and activity of repeated rescue sites, two key points emerge from our observations. First, rescue sites do not always promote repeated rescues. Out of all rescue sites that we observed, only 35% exhibited a second rescue when a subsequent catastrophe returned the depolymerizing end to that same position (Figure 4B). Furthermore, when we did observe two rescues at the same position, 43% of these microtubules went on to exhibit a third rescue while 57% were subsequently lost to depolymerization (data not shown). Second, rescue sites can promote repeated rescues over a relatively long period of time. We observed rescues occurring at a similar lattice site as soon as 30 seconds after a previous rescue, or up to 399 seconds later (data not shown). This indicates that sites can maintain rescue-promoting activity over a timespan that is much longer than the expected lifespan of GTP at the exchangeable site (i.e. ‘E-site’) of tubulin under our reaction conditions. In addition, we observed rescues occurring all along the length of the lattice, without bias toward the GTP-rich plus-end (Figure 5). These observations lead us to speculate that alternative lattice heterogeneities, independent of nucleotide state, could be required for rescue site activity. For example, previous work has shown that the microtubule lattice can transition between different numbers of protofilaments along its length, which could create a regions of lattice where protofilaments are ‘missing’ and the remaining protofilaments lack laterally bound neighbors^33^. We speculate that such an environment could disrupt the depolymerizing end by uncoupling the strain of outwardly curling protofilaments from subunits in the lattice (Figure 7).

Although our results are consistent with the presence of rescue sites in the lattice, we also show that the formation of these sites is not sufficient to explain rescue activity. Our wash-in experiments provide the best evidence that depolymerization conditions determine rescue activity more strongly than lattice polymerization conditions. By observing the same microtubules under alternating rescue-promoting or inhibiting conditions, we found that introducing rescue-promoting conditions during depolymerization is sufficient to recapitulate the rescue behavior of control microtubules that are constitutively exposed to rescue-promoting conditions (Figure 6). The key to these experiments was our finding that the divalent cations magnesium and calcium have potent effects on rescue activity. We speculate that high concentrations of divalent cations alter the structure of the depolymerizing end to inhibit rescue.

How could divalent cations inhibit rescue at the depolymerizing end? Divalent cations, like calcium and magnesium, are known to play important roles in microtubule dynamics. Magnesium promotes polymerization and depolymerization, and some of these roles are related to effects on GTP-binding and exchange ^34–36^. However, magnesium also binds tubulin at additional sites outside of the GTP-binding pockets ^37–40^. Hence, magnesium may have additional roles in regulating tubulin activity. We have shown here that both magnesium and relatively small concentrations of calcium (estimated 1nM free calcium; see Materials and Methods^41^) can drastically increase depolymerization rates, consistent with previous work ^25,29,42^. We also show that both cations are potent inhibitors of microtubule rescue (Figure 5). This suggests that the divalent cations may be suppressing rescue by altering how microtubules depolymerize. During depolymerization the protofilaments of a microtubule curl away from the lattice ^2^, which would bring the negatively charged regions on the outer surface of tubulin subunits into close proximity (Figure 7). Divalent cations could stabilize the outwardly curled conformation of the depolymerizing end by bridging the negatively charged side chains on the outer surface of tubulin (Figure 7). Consistent with this notion, we have shown previously that magnesium promotes depolymerization in a manner that requires the negatively charged c-terminal tail (CTT) domain of β-tubulin^25^. Interestingly, we observed that removing the β-CTT slows depolymerization and also promotes microtubule rescue, mimicking the effect of low magnesium (unpublished data;^25^). Therefore, divalent cations may inhibit rescue by binding the negatively charged CTTs and promoting a conformation of the depolymerizing end that favors depolymerization. Further studies are needed to determine how CTTs could change the activity or abundance of rescue sites in the lattice.

Our findings on the intrinsic rescue behavior of tubulin proteins could be extended to cells, where extrinsic proteins may further regulate rescue by binding and selectively stabilizing structural intermediates at the depolymerizing end and/or rescue sites in the lattice. For example, CLASP proteins promote rescue and suppress catastrophe, however, with only a modest effect on polymerization and depolymerization rates ^22^. CLASP is known to bind both at microtubule ends and along the lattice, raising the possibility that CLASP could regulate rescue by either promoting a conformational state of the depolymerizing end that favors rescue or by promoting the formation of rescue sites in the lattice ^9,19,22^. CLIP-170 proteins also promote rescue, and localize to both microtubule ends and to ‘GTP-islands’ in the microtubule lattice ^15–17^. Interestingly, the N-terminal microtubule-binding domain of CLIP-170 is sufficient to promote rescues *in vivo* ^16^. When combined with purified tubulin *in vitro*, the N-terminal domain promotes rescue and induces the formation of curved tubulin oligomers at microtubule ends and in solution, hinting that the rescue mechanism may be linked to the stabilization of specific protofilament structures ^17^. Further investigations of how these and other regulators that promote rescue could yield important insights into the structure of a rescuing microtubule end, the nature of rescue sites in the lattice, and how these may be related.

## Materials and Methods

Chemicals and reagents were from Fisher Scientific (Pittsburgh, PA) and Sigma-Aldrich (Saint Louis, MO), unless stated otherwise.

### *In vitro* Microtubule Dynamics Assays

Assays to measure microtubule dynamics by TIRF microscopy were based on previously described methods ^25,26^. Double-Cycled microtubule seeds were assembled by incubating 20 µM rhodamine tubulin (Cytoskeleton, Inc; Denver, CO) in BRB80 buffer (80 mM PIPES brought to pH6.9 with KOH, 1 mM ethylene glycol tetraacetic acid (EGTA), 1 mM MgCl_2_; minor pH adjustments were made with NaOH) with 1 mM GMPCPP at 37°C for 30 minutes. Sample was then centrifuged at 100,000 x g for 10 min at 30°C and the supernatant was removed. Pellet was suspended in 0.8x starting volume of ice cold BRB80 buffer to depolymerize labile microtubules. An additional 1 mM GMPCPP was added and microtubules were polymerized at 37°C for 30 min and pelleted again. The pellet was suspended in 0.8x starting volume warm BRB80. The reaction was then gently pipetted 8-10 times to shear the microtubules, aliquoted into 1 µL volumes, and either used immediately or snap frozen and stored at −80°C.

Imaging chambers were assembled using 22×22 mm and 18×18 mm coverslips. The coverslips were cleaned and silanized as previously described ^25,26^. The prepared glass coverslips were stored in desiccators at room temperature until used. The coverslips were mounted in a custom fabricated stage insert and sealed with melted strips of Parafilm. GMPCPP-stabilized microtubule seeds were affixed to coverslips using anti-rhodamine antibodies (Fisher Scientific, Cat# A-6397; diluted 1:50 in BRB80). Chambers were flushed with 1% Pluronic-F127 in BRB80 to prevent other proteins from adhering to the glass and equilibrated with an oxygen scavenging buffer (40 mM glucose, 1 mM Trolox, 64 nM Catalase, 250 nM Glucose Oxidase, 10 mg/ml Casein) prior to free tubulin addition. The imaging buffer consisting of unpolymerized tubulin (15-20% Hylite-488 labeled tubulin (Cytoskeleton, Inc.) and 80-85% unlabeled porcine brain tubulin), 5 mM MgCl_2_, 1 mM GTP, the oxygen scavenging buffer, and BRB80 to 50 µL volume was then flowed into the imaging chambers. The chamber was sealed with VALAP (1:1:1 Vaseline, Lanolin, Paraffin wax) and warmed to 37°C using an ASI400 Air Stream Stage Incubator (Nevtek; Williamsville, VA) for 5 minutes before imaging. Temperature was verified using an infrared thermometer.

For experiments testing the impact of calcium on rescue activity, 10µM CaCl_2_ was added to the imaging buffer prior to chamber assembly. We used MaxChelator (maxchelator.stanford.edu) to estimate the concentration of free calcium in these reactions using theoretical values for ionic and chelator concentrations, pH and temperature. At standard conditions plus calcium (5 mM MgCl_2_, 10 µM CaCl_2_, 1 mM EGTA and 1 mM GTP at pH:6.9 and 37°C) it is estimated that more than 99.9% of the Ca2+ is bound to EGTA.

Images were collected on a Nikon Ti-E microscope equipped with a 1.49 NA 100× CFI160 Apochromat objective, TIRF illuminator, OBIS 488-nm and Sapphire 561-nm lasers (Coherent; Santa Clara, CA), and an ORCA-Flash 4.0 LT sCMOS camera (Hammamatsu Photonics; Japan), using NIS Elements software (Nikon; Minato, Tokyo, Japan). Images were acquired using two-channel, single-plane TIRF, at 1 second intervals.

### Image Analysis

Images were analyzed using a custom-made MATLAB program as described previously^25^. Briefly, seeds were identified by thresholding image intensity, and used to segment the images along the axis of the microtubule. Images were then automatically cropped to 4 pixels above and below the microtubule axis, then the max intensity projection for each timepoint were stacked to generate kymographs for analysis.

Polymerization and depolymerization rates were calculated by measuring the changes in microtubule length and time from the first and last points of the individual polymerization and depolymerization events from the kymographs. Polymerization rate constants were estimated as the slope of the polymerization rate linear model. Depolymerization rate constants were determined similarly but using the median depolymerization rate (µm*min^−1^) from all tubulin concentrations pooled.

Rescue and catastrophe frequencies were calculated as the quotient of the number of transitions and depolymerization or polymerization duration respectively. Rescue abundance was determined by dividing the number of unique rescue events by the total length lost during depolymerization. Unique rescue events were defined as rescue sites separated by at least 200 nm from previous rescue sites on the microtubule lattice. Rescue position relative to the plus- end was calculated as the difference between microtubule length at catastrophe and length at rescue.

### Tubulin wash-in experiments

For wash-in experiments, GMPCPP-seeded imaging chambers were similarly assembled, but not sealed with VALAP. Imaging chambers were warmed on the stage for 3-5 minutes, allowing the temperature to equilibrate to 37°C, then dynamic microtubules were imaged for 10 minutes before the imaging chamber with was flushed 4-5x chamber volumes of warm reaction buffer. Image acquisition was paused during the chamber flush and resumed immediately afterwards. The average wash-in duration for all wash-in experiments was ∼60 seconds from time of acquisition pause to resumption.

Images were processed as described above, with the addition of post-acquisition image stabilization that was used to reduce minor XY drift during image acquisition using the Image Stabilizer Plugin for ImageJ ^43^.

### Determining Tubulin Concentration

For each experiment, the tubulin concentration was determined by running a sample of the imaging buffer containing unpolymerized tubulin on a 10% Bis-Tris SDS-PAGE gel, followed by staining with Coomassie blue. The concentration was determined by densitometry, directly compared to standard curve made from BSA standards run on the same gel.

### Monte Carlo Simulations

Microtubule dynamics were simulated using a custom-made MATLAB program. The simulation modeled microtubule length changes over time based on empirically derived values for polymerization and depolymerization rates as well as transition frequencies. Rescues were simulated slightly differently between the two models; either stochastically based on the experimental rescue frequency for the free model. Rescues were simulated as sites incorporated into the growing microtubule at an empirically derived frequency. Following catastrophe, a rescue event was triggered as the depolymerizing end approached the incorporation site within 100 nm. For simplicity, rescue sites were lost after activation, resulting in only a single rescue per site. The simulated microtubules lengths were analyzed as experimental data.

## Acknowledgements

We thank Dr. Melissa Gardner (University of Minnesota) for technical assistance in developing *in vitro* assays to measure microtubule dynamics, Dr. Jay Gatlin (University of Wyoming) for helping with the purification of porcine brain tubulin, and members of the Moore lab for helpful discussions and advice. This work was supported by National Institutes of Health (NIH) grant R01GM112893 (to J.K.M.).

## Author Contributions

C. Fees: Conceptualization, data curation, formal analysis, validation, investigation, methodology, resources, software, visualization, and writing – original draft, reviewing and editing.

J. Moore: Conceptualization, resources, supervision, funding acquisition, investigation, methodology, visualization, and writing – original draft, reviewing and editing.

---

**Supplemental Figure 1)**

**(A)** Percent of seeds with at least one observed rescue as a function of tubulin concentration. Protein purified from pig brain (black) compared to purchased protein purified commercially (red). Data points represent mean ± 95% CI of 870 separate GMPCPP seeds from 8 different experiments. **(B)** Average number of rescues per seed as a function of tubulin concentration.

**Supplemental Figure 2)**

**(A)** Polymerization rate as a function of tubulin concentration of reactions with (red) or without (black) 10 µM CaCl_2_. Points represent average microtubule polymerization rate at each concentration.

**Supplemental Figure 3)**

**(A)** Polymerization rate constant of each MgCl_2_ concentration determined as the slope of the regression of polymerization rate as a function of tubulin concentration. **(B)** Depolymerization rates of microtubules that rescue (solid bars) and microtubules that do not rescue (empty bars). Bars represent at least 283 microtubules from at least 3 separate experiments for each condition.

